# Methods for computing the maximum performance of computational models of fMRI responses

**DOI:** 10.1101/377101

**Authors:** Agustin Lage-Castellanos, Giancarlo Valente, Elia Formisano, Federico De Martino

## Abstract

Computational neuroimaging methods aim to predict brain responses (measured e.g. with functional magnetic resonance imaging [fMRI]) on the basis of stimulus features obtained through computational models. The accuracy of such prediction is used as an indicator of how well the model describes the computations underlying the brain function that is being considered. However, the prediction accuracy is bounded by the proportion of the variance of the brain response which is related to the measurement noise and not with the stimuli (or cognitive functions). The bound to the performance of a computational model to the prediction of brain responses has been referred to as the noise ceiling. In previous neuroimaging applications two methods have been proposed for estimating the noise ceiling based on either a split-half procedure or Monte Carlo simulations. These methods make different assumptions over the nature of the effects underlying the data, and, importantly, their relation has not been clarified yet. Here, we use a two-level generative framework to formally describe the partition between the variance of measurement noise and the stimulus related variance. In this framework we derive an analytical form for the noise ceiling that does not require computationally expensive simulations or a splitting procedure that reduce the amount of data. We describe the relation between the newly introduced noise ceiling estimator and the previous methods for variable levels of measurements noise using simulated data. Additionally, as the relation to the noise ceiling is used to make conclusions on the validity of a model with respect to others, we evaluate the effect the interplay between regularization (often used to estimate model fits to the data when the number of computational features in the model is large) and model complexity on the performance with respect to the noise ceiling. Finally, we show the differences between the methods on real fMRI data acquired at 7 Tesla. We demonstrate that while the split half estimator provides a pessimistic estimate of the noise ceiling due to the small amount of data available in conventional fMRI datasets, the parametric nature of the Monte Carlo estimator results in overly optimistic estimates. For this reason, for real data, we propose a robust procedure to the estimation of the noise ceiling based on bootstraps.

**Author Summary:** Encoding computational models in brain responses measured with fMRI allows testing the algorithmic representations carried out by the neural population within voxels. The accuracy of a model in predicting new responses is used as a measure of the brain validity of this model, but the result of this analysis is determined not only by how precisely the model describes the responses but also by the quality of the data. In this article, we validate existing approaches to estimate the best possible accuracy that any computational model can achieve conditioned to the amount of measurement noise that is present in the experimental data (i.e. the noise ceiling). Additionally we introduce a close form estimation of the noise ceiling that does not require computationally or data expensive procedures. All the methods are compared using simulated and real fMRI data. We draw conclusions over the impact of regularisation procedures and model complexity and make practical recommendations on how to report the results of computational models in neuroimaging.

## Introduction

Computational modelling approaches applied to functional magnetic resonance imaging (fMRI) measurements aim to explain and predict (patterns of) brain responses by expressing them as a function of model features that describe the sensory (or cognitive) stimuli [1–5]. By doing so, computational neuroimaging methods have been proposed as a means to test the (brain) validity of the algorithm being evaluated and eventually its refinement.

At the single voxel level, two different approaches, population Receptive Fields (pRF) modelling [3] and linearized encoding models [6, 7], have been developed to link computational models and fMRI responses. In the following, we will refer to both these approaches indiscriminately as encoding models (see e.g. [8] for the relation between linearized encoding models and pRF approaches). The performance of a computational model that describes fMRI responses is (most often) evaluated in terms of its accuracy in predicting new (test) data. An issue that presents itself when evaluating modelling efforts (often across laboratories and data sets) is that prediction accuracy is not only affected by inaccuracies in the definition of the algorithm (i.e. mismodelling) but also by other sources of variance in the brain responses that are not expressly modelled (e.g. attention and adaptation) and, most importantly, by physiological (e.g. respiration) and measurement noise. These effects are evidenced by the fact that, in real data, presenting multiple times the same stimulus does not result in the same measured brain response. Commonly tested models of sensory (or cognitive) stimuli do not account for the variability in the response between repetitions of the same stimulus which imposes a bound to the ability to encode computational models in fMRI responses. This bound can be interpreted as the performance of the computational model underlying the generation of the responses (i.e. the true underlying model) given the noise (experimental, physiological or other) that is present in the data and has been referred to as the *noise ceiling* [6,9–11]. In the neuroimaging community, it has been recommended to report the performance of a computational model with respect to the noise ceiling and, in some cases, these recommendations have led to the use of normalized accuracy scores (e.g. dividing the accuracy by the noise ceiling, [11–15].

To illustrate the concept of noise ceiling we can consider the process of fitting a computational model using an encoding approach. Considering an fMRI experiment in which brain responses are measured to the presentation of several sensory stimuli (each one repeated multiple times), in the perspective of hierarchical statistical models, the encoding procedure is equivalent to estimating the brain response to each stimulus from the fMRI time series at the first level and, at the second level, estimating how these responses can be explained by a computational model at hand. A consequence of this hierarchical framework is that the prediction accuracy at the second level is bound by the uncertainty in the estimation of the evoked response at the first level. The value of this bound corresponds to the intraclass correlation coefficient, a well-known result in multilevel modelling. While this two-level procedure is not used by all encoding approaches in practice, the unmodelled variability of the response between repetitions of the same stimulus limits the performance of a computational model that predicts the whole fMRI time series as well (see e.g. pRF models [3]). Reporting the noise ceiling allows assessing the quality of the predictions obtained when using computational modelling approaches relative to the quality of the data, and thus comparing modelling efforts on different datasets across labs.

The noise ceiling can be obtained considering the variability across subjects (see e.g. [9]), but here we will focus on estimation procedures at the single subject level, where two approaches have been proposed to estimate the noise ceiling of single voxels. The first, models the response of a voxel as a univariate normal distribution with two variance components [16]. The first variance component corresponds to the variability of the signal around its mean due to genuine differences in the brain response between different stimuli (excluding the effects of measurement noise). The second variance component corresponds to the variability in the brain response due to measurement noise. Having an estimate of the measurement noise allows generating new samples for both the signal without noise (genuine brain response) and the measurement (i.e. signal plus noise) using Monte Carlo simulations. The noise ceiling (measured with correlation or predictive R^2^) is then computed using the simulated signals and measurements (i.e. considering the performance in predicting the noisy measurements of a model whose prediction is the clean signal). In what follows we will refer to this approach as to the Monte Carlo noise ceiling (MCnc). Alternatively, the noise ceiling can be estimated as the correlation between the estimates of the responses in two independent repetitions of the same experimental procedure [17, 18]. In absence of two repetitions of the test set, the split-half noise ceiling estimator (SHnc) can be estimated by splitting the available test data in two disjoint sets (i.e. splitting the trials of all test stimuli in two sets so to obtain two estimates of the test data), computing the split-half correlation and applying a correction factor that accounts for the reduced number of trials in each half of the data compared to the full dataset.

In this article, we describe the differences between these two noise ceiling estimators using data simulated using a generative framework. We derive an analytical solution to the calculation of the noise ceiling obviating the need of computationally demanding procedures (Monte Carlo simulations) or splitting the data in two sets. The validity of the analytical solution is also tested using simulated data. When using linearized encoding approaches, regularization is often required because of the dimensionality of the model with respect to the number of stimuli and because of collinearity between the features of the computational model. Here we evaluate how the bias variance tradeoff introduced by the regularization influences the performance of an ideal model and thus the relationship to the noise ceiling (which is model independent) by imposing a second constraint.

Finally, we evaluate the differences between the noise ceiling estimators using real fMRI data, obtained from 7 Tesla acquisitions, and considering both cortical and sub-cortical regions. Both the MCnc and the analytical noise ceiling depend on the estimation of the variance of the brain responses. For this reason, in the analysis of real data, we evaluate the impact of using non-parametric estimates (using bootstrap) of this variance on the calculation of the MCnc and analytical noise ceiling. We demonstrate that the previously used parametric estimate leads to an over estimation of the noise ceiling. Our results are discussed in terms of their implications for the evaluation of computational models and their comparison using fMRI data.

## Methods

We consider a two-level procedure to fit a computational model to fMRI responses. At the first level, fMRI responses to single stimuli are estimated from the fMRI time series and, at the second level, the computational model is (linearly) fit to the vector of fMRI responses in order to derive the parameter weights (in the space of the features of the computational model – the population receptive field of a linearized encoding model). As highlighted in the introduction, while alternative approaches exist that estimate and predict the fMRI signal at the level of the whole time course or use non-linear models [3, 19], the target of computational models of the fMRI response is to predict the voxels’ responses to single stimuli and the two-level procedure allows formalizing a generative framework for (linearized) encoding models and introducing the notion of noise ceiling in relation to known properties of hierarchical statistical models.

Fig 1 illustrates key concepts that underlie the notion of noise ceiling. In a generative framework (blue rectangle in Fig 1) brain responses can be considered as a function of the representation of the stimuli in the space of a computational model (3-features in Fig 1). Nevertheless, the measurement of brain responses is affected by experimental/measurement noise such that the estimate of the responses of a given voxel to the same stimuli in two independent measurements (fMRI run1 and 2 in Fig 1) differs. Fitting a computational model to the measured brain responses allows linking the model features to the measurements and thus predicting voxel responses to new (test) stimuli. When this procedure is used on noisy measurements, the experimental noise imposes a limit to the performance of the model in predicting the measured responses.

**Fig 1.**
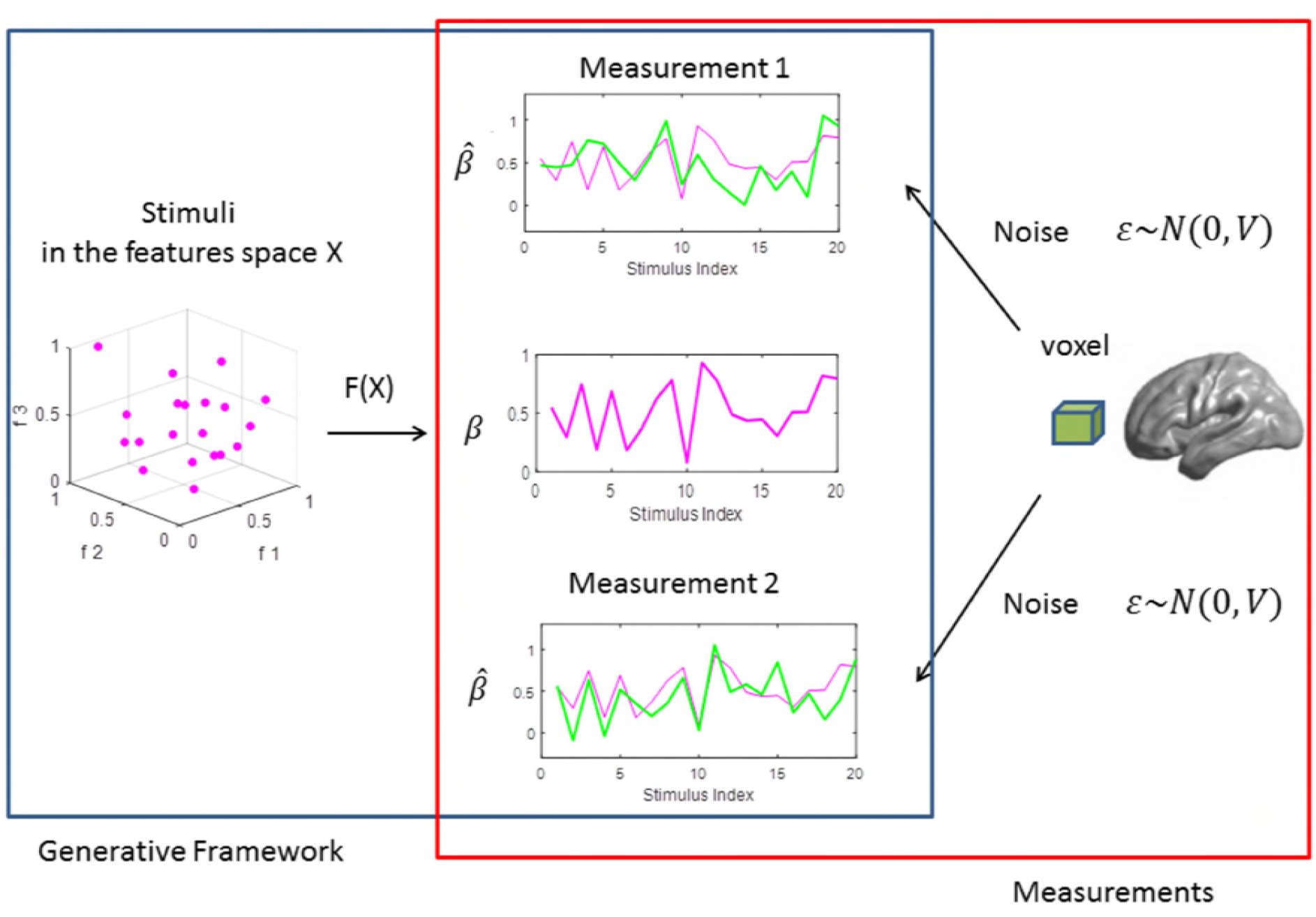
General description of the relation between the generative framework (left to right) and the estimation framework (right to left). At the generative level, brain responses (β – pink curve) are obtained as a function of the model based representation of the stimuli (pink dots). The experimental noise corrupts the observation of the brain responses to the stimuli resulting in two different measurements (green curves) in the two disjoint sets (fMRI run1 and run2).

In what follows we first describe the two-level fitting approach and the different metrics used to evaluate model fitting in order to introduce some relevant concepts, next we introduce a generative framework and derive the bound to the performance.

### Estimation of the response to single stimuli

At the first level, the observed fMRI response is assumed to be linearly dependent on the stimuli (design matrix) [20, 21] and the estimation (for every voxel) is achieved using e.g. Ordinary Least Squares (OLS):

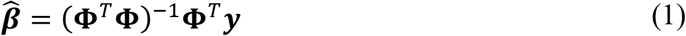

For every voxel, 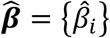 is the vector of the estimated responses to the *n* stimuli, ***y*** is vector of the voxel time course (the observed fMRI signal) and **Φ** is the design matrix describing the timing of presentation of the stimuli in the experiment, including the effect of the hemodynamic response. Under the assumption of identically independent distributed errors and a full rank design matrix the estimated responses 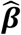 are normally distributed around their expected value with covariance equal to 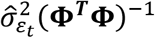, where and 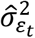 is the estimated variance of the experimental noise of the voxel time course [21].

Note that violations of the OLS assumptions will produce biased estimates of the 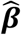 variances, even in the cases where the estimation of the 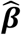 is unbiased [22]. In this sense, the presence of temporally auto-correlated noise in the fMRI time series requires the use of Generalized Least Squares (GLS) which is based on an estimation of the structure of the temporal correlations [23, 24]. The 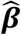 variances estimated with OLS can also be biased due to the presence of non-stationary noise or due to the random effects of the brain response across e.g. the fMRI runs/sessions [25]. In order to obtain reliable estimators 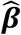 variances non parametric methods [26] or mixed effects models can be used as an alternative to the OLS/GLS estimators.

### Linearized encoding models

Linearized encoding models assume the estimated response vector 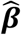 to be linearly dependent on the description of the stimuli on the basis of a computational model represented by a matrix ***X*** that projects each of the *n* stimuli onto the model space described by a model with *f* features (Fig 1). The objective of encoding approaches is to model the differences in brain responses between stimuli (at each voxel) as a function of the representation of the stimuli in the feature space. As a consequence, the mean of the brain response at each voxels is not interesting and should be removed from the response vector 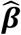 before fitting the model [27]. When there is collinearity across the features or when the number of stimuli *n* is smaller than then number of features *f*, regularization is used. We consider here the use of regularization as in fMRI it is common for *n* to be smaller than *f* and we want to evaluate the impact of regularization in comparing the performance of a model to the noise ceiling. Without loss of generality we consider the use of ridge regression [28] as a form of regularization. For every voxel, the estimate of the linear weights 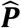 that link the computational model (represented by the matrix ***X***) to the estimated voxel response vector 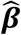 is:

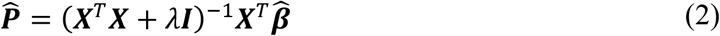

Where *λ* is the regularization parameter, 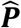 represents a (linearized) estimate of the voxel pRF and allows predicting the responses to new (Test) stimuli by considering 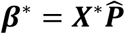, where ***X***\* contains the representation of the stimuli in the test set in the space of *f* features. Other approaches use grid search or more sophisticated optimization algorithms when a non-linear relationship between the features and the response is assumed [3, 19].

### Evaluating the performance of computational model encoded in fMRI responses

The performance of a computational model can be assessed on test stimuli using e.g. the sample correlation coefficient between the responses predicted by the computational model 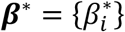 for every stimulus *i*, and the estimated responses with OLS/GLS at first level 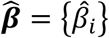 (see Eq. 1):

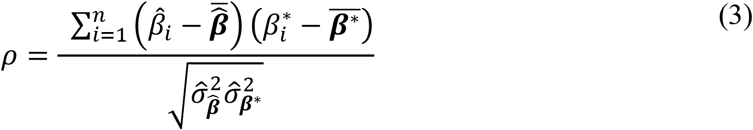

where the variances refer to the variability between the components of the vector of responses around the mean of the vector: 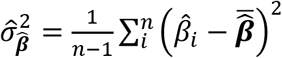. Alternatively, predictive *R*^2^ is frequently used for describing the performance of an encoding model:

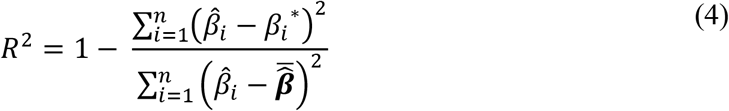

where 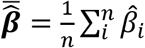, is the mean of the estimated response across its components (each component corresponds to one presented stimuli). Here we are computing the explained variance between the observed and predicted brain responses 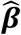 and ***β***\*, which is different than the explained variance obtained with OLS/GLS in the domain of the fMRI time series ***y***. When OLS is used and the performance is computed with the training data, the value of *R*^2^ is contained in the interval [0, 1], however when *R*^2^ is computed on an independent (test) data set this interval is not valid and any number value in the interval [–∞, 1] can occur. It is important to note that, while *R*^2^ is sensitive to scaling transformations of the estimated response, the correlation coefficient measures the similarity between the predicted and observed responses in term of covariations around their mean and is insensitive to scaling transformations. This difference between the metrics is relevant when regularization is used. The relation between predictive *R*^2^ and *ρ* can be rendered explicit considering (without loss of generality) the estimated responses were centred to have zero mean 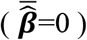 [27]:

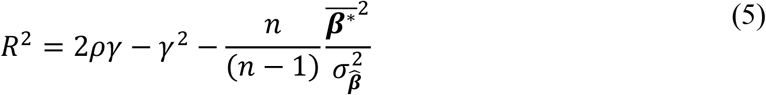

Where *γ*^2^ is the ratio between the estimated variances of the observed and predicted response vectors and 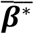 represents the bias in the mean of the predicted response vector (i.e. how much the mean of the predicted response vector differs from zero, Appendix I). The Euclidean distance *D*^2^ between the vectors 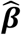 and *β*\*, which is also a frequently used metric is closely related to the explained variance by: 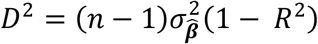.

### Generative framework for linearized encoding

We consider a generative framework that matches the assumptions of the linearized estimation procedure highlighted above. The estimated response vector 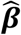 is assumed to be multivariate normally distributed around the mean vector ***β*** and with covariance matrix that reflects the experimental variability of the 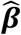 estimator (i.e. the experimental noise of the fMRI data represented by the estimated variance of the residuals in the time domain 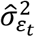, and correlations imposed by the design matrix Φ). Note that this can include errors introduced by misspecification of the design matrix (e.g. an erroneous definition of the hemodynamic response). For OLS and GLS estimators the 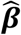 variance-covariance matrix takes the form of:

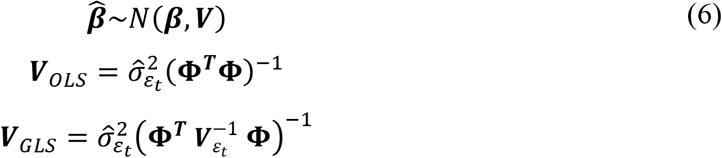

Where ***V**_ε_t__* defines the covariance matrix of the fMRI noise in the time domain, typically estimated with autoregressive models [22]. When considering linearized encoding models, ***β*** (i.e. the mean of the estimated response) can be considered to be generated on the basis of the computational model defined by the matrix ***X***. In particular we can consider:

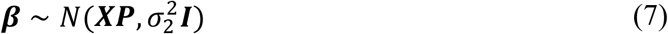

At this second level the noise variance 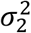 describes only mismodelling effects at the level of the computational model (i.e the effect of features not comprised by ***X***) and ***P*** is the population receptive field corresponding to the features contained in ***X***.

Note that this generative framework assumes a multivariate normal distribution for 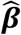. As a consequence each component of 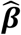 (the response to each stimulus) can have different variance. Considering this generative framework, when we predict the response vector of new test stimuli using a linearized encoding approach, the variance of the estimated response vector can be partitioned in two components, the first reflecting the measurement variability and the second describing the validity of the encoding model.

### Methods for estimation of the performance of linearized encoding models Split-half noise ceiling (SHnc)

The split-half noise ceiling estimator is an empirical procedure which consists of correlating the 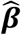 responses to the same stimuli obtained from two disjoint sets of data: (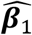 and 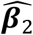) at every voxel [17,29,30]. Because the number of stimuli at each split is reduced by half, the observed correlations are adjusted using the Spearman-Brown correction. The split-half noise ceiling is a non-parametric procedure since it does not rely on the estimation of the 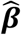 variances. Therefore, the split-half noise ceiling estimator procedure takes into account all sources of variability that affect the brain responses:

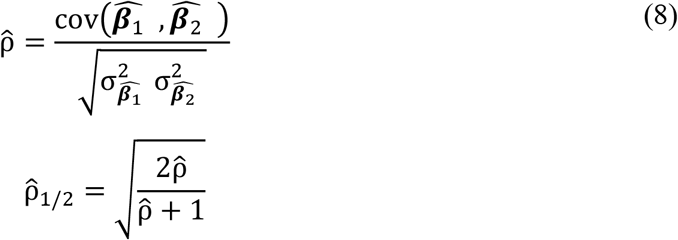

Note that the SPnc is defined only for positive split half correlation values. Here we define the SPnc to be zero for negative split half correlations, which is equivalent to assuming that in the case in which the observed correlation between two independent measurements of the same stimuli is negative the maximum performance that any encoding model can achieve is the chance level.

### Monte Carlo noise ceiling (MCnc)

The MCnc assumes every component of the response vector (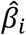, with *i* running across stimuli) to be samples of the same univariate normal distribution [16]. The variance of this distribution is assumed to be formed by two components, the experimental noise 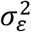 and the variability between stimuli of the *noise free* ***β*** responses 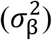.

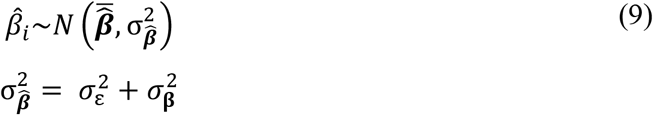

Here we make the distinction between 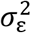 and the 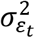. (in Eq. 6), 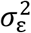 refers to the average variability in the estimation of ***β*** across its components (pooled), while the 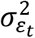. in refers to the variability of the residuals of the domain of the fMRI time series (Eq. 1). The variance of the noise 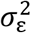 in Eq. 9 is obtained considering the variance of the estimated responses pooled across stimuli (see Appendix II). The variance of the response of each stimulus (i.e. each component of 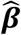) can be obtained with OLS or with GLS. When pooling across stimuli the MCnc assumes that the noise affects all stimuli in the same way. The variability of the *noise free* signal is obtained as the difference between the variability of the 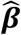 responses across the *n* stimuli 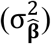 and the variability of the noise:

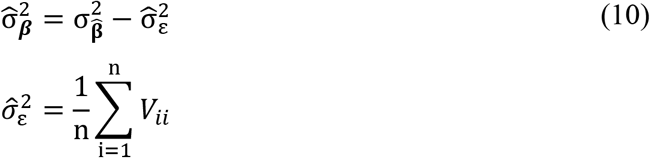

The term 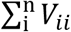 corresponds to the sum of the diagonal entries of the 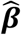 covariance matrix, derived from the OLS/GLS estimation of the 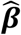 coefficients. After the variability of the *noise free* signal is obtained with Eq. 10, Monte Carlo samples of the *noise free* signal can be generated. For every generated noise free signal the experimental noise is added (using a normal distribution with zero mean and variance 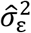) and the correlation between the *noise free* signal and the *noise contaminated* signal is computed. The median across all simulations represents an estimate of the noise ceiling.

### Analytical derivation of noise ceiling (NC)

Following the two-level generative model described in Eq. 6,7 the noise ceiling is defined as the expected performance (measured as correlation or predictive *R*^2^) of the model underlying the generation of the responses (i.e. the “true” model). This model uses the generative pRF (***P***) and produces correct predictions without the effects of neither measurements nor mismodelling errors, which in practice means: ***β* = β:.= XP*** and 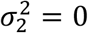, see Eq. 7. According to this definition the noise ceiling for the correlation coefficient is:

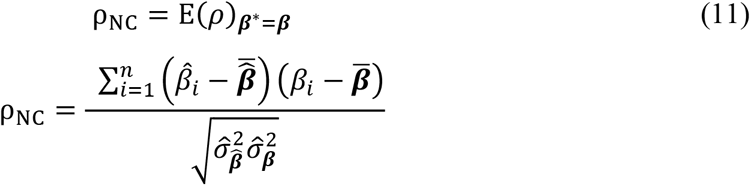

In the framework of multi-level models, the noise ceiling is equal to the intraclass correlation coefficient [31], which is defined as the ratio between the standard deviations of level 1 (Eq. 6) and level 2 (Eq. 7), (see Appendix II):

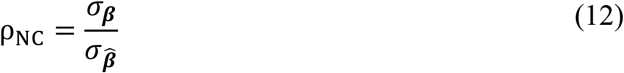

The denominator 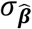 is easy to estimate since the values of 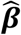 are known, however the numerator *σ*_***β***_ should be estimated using the covariance matrix of the estimated 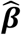. The analytical noise ceiling can be estimated using the variance estimates of each component of 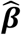 derived from OLS/GLS, according to the formula:

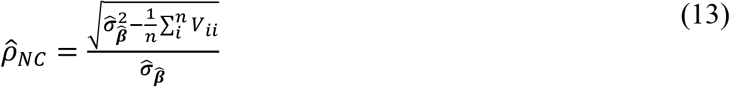

Note that in Eq. 13 a negative number under the square root can be obtained if the average of the variance within components (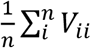 is greater that the variance between components 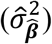. This can be caused by one or more stimuli (components of 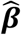) with a brain response that is equal to zero. In these cases we define the analytical noise ceiling to be zero, which corresponds to assume that the maximum accuracy that encoding models can reach is the chance level. The same indeterminacy of obtaining a negative estimation of the variance can occur for the Monte Carlo NC (Eq. 10) and the considerations made for the MCnc and the analytical noise ceiling are consistent with the ones made for the SHnc in the case of negative correlations between the splits. In the case that the assumptions of the OLS/GLS model are suspected not to be valid a non-parametric estimator of the variance of the components of the 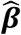 can be used (see real data applications in the result section).

The noise ceiling estimator 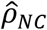 has as expected value ρ_NC_, (see appendix II for the demonstration). The noise ceiling estimator for the explained variance R^2^ can be derived transforming the noise ceiling for the correlation coefficient to the R^2^ domain with Eq. 5 or analytically from the definitions in the generative framework. Both approaches produce the same result of: 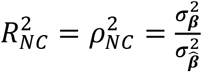. The analytical noise ceiling estimator has the same expected value than the Monte Carlo noise ceiling estimator with the advantage of not requiring a large computational effort.

### Simulations

Simulated fMRI time series were obtained, following our generative approach, starting from a computational model (***X*** in 128 features) and assuming a linear encoding model where the responses to the stimuli ***β*** are obtained from a linear combination (the population receptive field) of the features of the model. The simulated population receptive field vector ***P*** was sampled from the standard normal distribution. To simulate mismodelling (when required) we added to the simulated responses gaussian random noise with variance 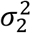 of different values according to Eq. 7. The ***β*** vector was then combined with a plausible design matrix ***φ*** describing the presentation of the stimuli during an fMRI experiment. In particular the design followed a fast event related design identical to the one used in the acquisition of the real data (see below). Experimental noise was simulated considering additive Gaussian random noise (with variance 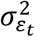 according to Eq. 6) at the level of the generation of the simulated fMRI time series. Each simulation was replicated 1000 times under identical conditions and we report the average value across the simulations and the [5,9,5] percentiles of the distribution. To avoid the variance of the 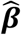 is to be affected by adding mismodelling effects (when required), the vector ***β*** was standardized such that 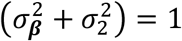 and 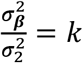. As result, the mismodelling effects are expressed in relative terms with respect to the variance of the model 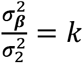.

### MRI data

The data presented here are part of a larger study that includes N=10 healthy participants. The subjects had no history of neurological disease, and gave informed consent before commencement of the measurements. The Ethical Committee of the Faculty for Psychology and Neuroscience at Maastricht University granted approval for the study. Magnetic resonance imaging data were acquired on an actively shielded MAGNETOM 7T whole body system driven by a Siemens console at Scannexus (www.scannexus.nl). A Nova Medical head RF coil (single transmit, 32 receive channels) was used to acquire anatomical (T_1_, Proton Density [PD] weighted) and functional (T_2_* weighted BOLD) images. T_1_ weighted (0.7 mm isotropic) images were acquired using an MPRAGE sequence (repetition time [TR] = 3100 ms; time to inversion [TI] = 1500 ms; time echo [TE] = 3.5 ms; flip angle = 5°). PD images were acquired with the same MPRAGE as the T1 weighted image but without the inversion pulse (TR = 2160 ms; TE = 3.5 ms; flip angle = 5°), and were used to minimize inhomogeneities in T_1_ weighted images [32]. Acquisition time for the T_1_ and PD datasets were ~ 9 and 4 minutes respectively. Anatomical data were analyzed with BrainVoyager QX and were resampled (with sinc interpolation) in the normalized Talairach space (Talairach and Tournoux, 1988) at a resolution of 0.5 mm isotropic.

Functional (T_2_* weighted) data were acquired using a clustered Echo Planar Imaging (EPI) technique (1.1 mm isotropic; TR = 2.6 s; GRAPPA = 3; MultiBand = 2; Gap = 1.4 s). The experiments were designed according to a fast event-related scheme ad slices were prescribed in a coronal oblique orientation in order to cover the brainstem and auditory cortex (Heschl’s gyrus, planum temporale and planum polare) bilaterally. A total of 168 sounds were presented six times across 24 runs in silent gaps in between volme acquisitions using magnetic compatible earbuds (Sensimetrics inc.). The sounds were divided into four training and testing sets (126 and 42 sounds respectively). Within each run, sounds were randomly spaced at a jittered interstimulus interval of 2, 3, or 4 TRs and presented in the middle of the silent gap between acquisitions (leaving 100 ms of silence before and after the sound). Zero trials (trials where no sound was presented [5% of the trials]), and target trials (trials in which a sound was presented two times in a row [5% of the trials]) were included. Subjects were instructed to perform a one-back task, and were required to respond with a button press when the same sound was presented two times consecutively. Target trials were excluded from the analysis. Before starting the experiment (with the ear buds in place), the subjects were instructed to adjust the overall sound intensity to a clearly audible and comfortable level. This resulted in an approximate sound intensity of 65 dB. The total scanning was divided over two sessions that were acquired in two consecutive days.

### Functional data analysis

Functional data were analyzed with BrainVoyager QX [33]. Preprocessing consisted of slice scan-time correction (with sinc interpolation), 3-dimensional motion correction, and temporal high pass filtering (removing drifts of 4 cycles or less per run). Functional data were coregistered to the anatomical data, normalized in Talairach space (Talairach and Tournoux, 1988), and resampled (with sinc interpolation) at a resolution of 1 mm isotropic (for the cortical analysis) and 0.5 mm isotropic (for the sub-cortical analysis). An additional spatial smoothing using a gaussian kernel with a standard deviation of 1 mm was applied to the sub-cortical data. We calculated the fMRI response to each sound in three steps. First, we computed noise regressors to denoise the data ([16], http://kendrickkay.net/GLMdenoise/). These regressors were added to the second and third step, which were otherwise executed as described before (see the analysis described in [34]). That is, as a second step an optimized hemodynamic response function (HRF) per voxel but identical across sounds was computed using a deconvolution analysis. Third, the estimated HRF per voxel was used to estimate the amplitude of the response (i.e. beta weight) to each sound using ordinary least squares with a design matrix with as many predictors as sounds. Note that these steps were performed independently for training and testing runs. To avoid overfitting, the HRF estimation procedure was performed only on the training data and, for each voxel, the HRF estimated in training was used to estimate the test responses.

## Results

### Validating the analytical noise ceiling estimator

We validated the analytical noise ceiling estimator by comparing (mean and [5 95] percentiles across 1000 simulations) the estimated noise ceiling value (Eq. 13) with the performance of the model used for the generation of the data in simulations. This model predicts the brain responses using the generative ***P*** and ***X*** with 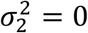 in Eq. 7.

Fig 2 shows the results of the comparison for both correlation and predictive R^2^. The mean value and the variance of the analytical NC estimator matched the mean value and variance of the performance of the generative model. The R^2^ showed stronger dependence with the experimental noise than the correlation coefficient.

**Fig 2.**
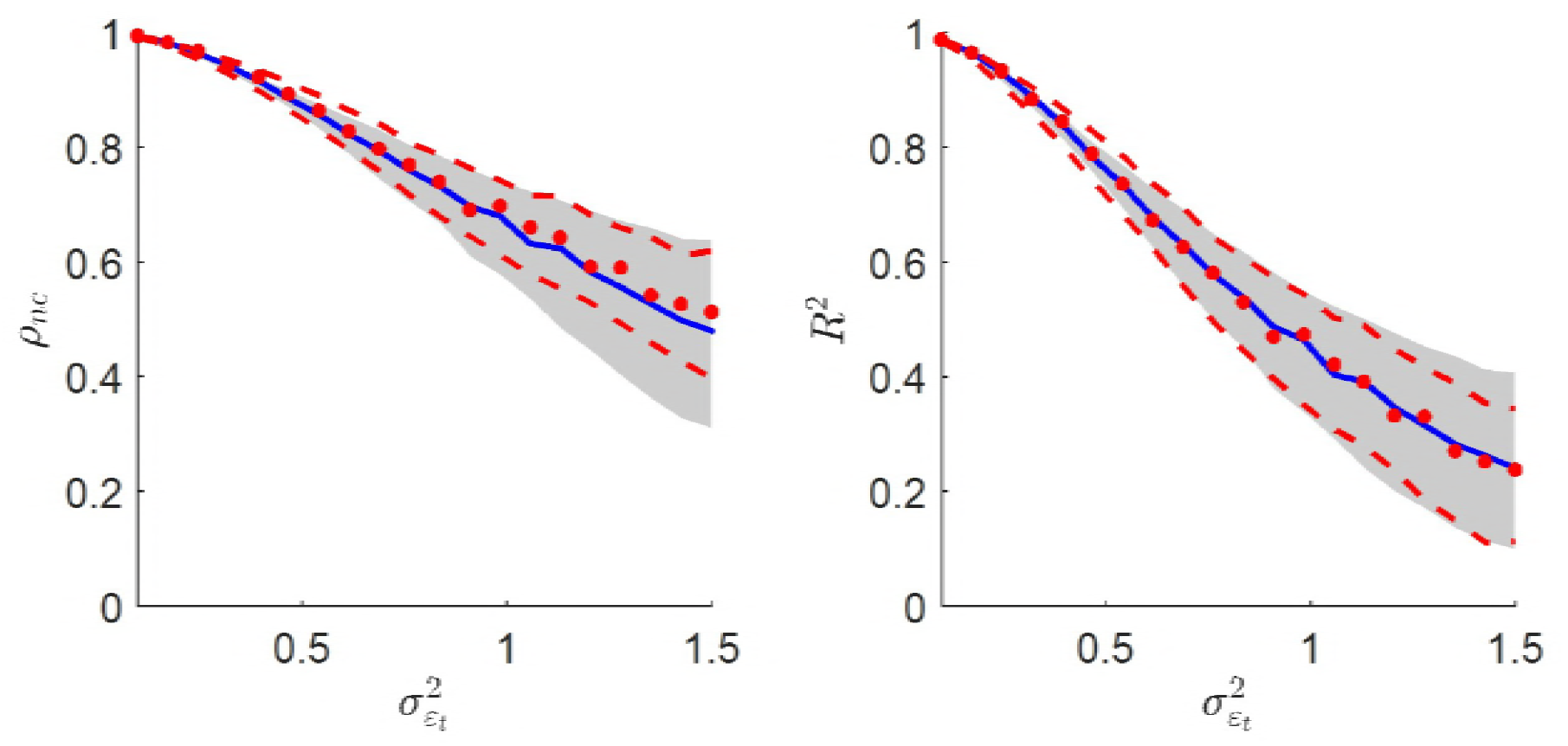
The analytical noise ceiling estimator (blue) and its true value obtained from the generative model (red dots) is presented for different levels of experimental noise 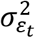. The left panel shows the noise ceiling for the correlation coefficient while the right panel shows the noise ceiling for the explained variance. The grey shadowed area denotes the [5 95] percentiles of the distribution, while the red dotted line denotes the [5 95] percentiles for the true value across 1000 simulations under the same generative parameters.

### Comparison with other methods

Figure 3 (left panel) shows the mean and [5 95] percentiles bands of the split-half estimator [17], the Monte Carlo estimator [16] and the analytical estimator (Eq. 13). We report only the results using the correlation as evaluation metric for simplicity. The three NC estimators resulted in the same mean estimate across the 1000 simulations. The SHnc estimates were characterized by wider confidence bands than the ones obtained with the MCnc and the analytical NC estimators. The MCnc and the analytical NC were characterized by the same variability (the [5 95] confidence bands are superimposed in Fig 3 left panel). For the three NC methods, the variability of the estimated NC increased with decreasing mean value (mean and variance are not independent). The relation between the three noise estimators is presented as a scatter plot in the right panel of Fig 3. The three estimators produced similar values when the noise variance is low, however in noisy scenarios the increase in the variability of the estimates produced large differences between the SHnc and the analytical noise ceiling. The MCnc and the analytical noise ceiling resulted in very similar estimates independent of the level of experimental noise. The advantage of using the analytical noise ceiling estimator (Eq. 13) is that it did not require a computationally expensive resampling procedure at each voxel.

**Fig 3.**
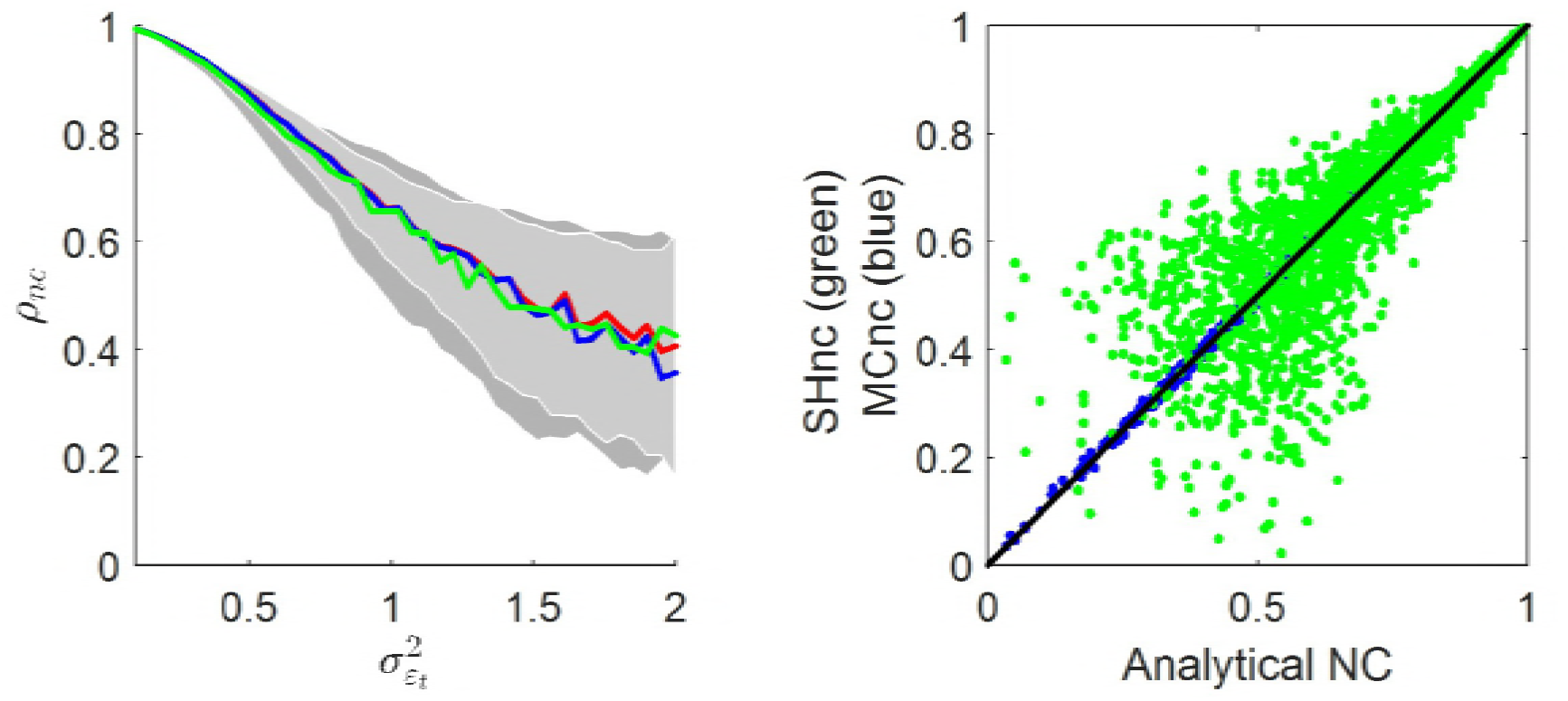
Left Panel: three noise ceiling estimators are presented as a function of the experimental noise 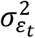. The mean of the NC across the 1000 simulations are presented in blue and red for the analytical and the Monte Carlo NC respectively. The mean of the split-half NC is presented in green. The darker shadowed area corresponds to the [5 95] percentiles of the distribution of the split-half estimator across the 1000 replications. The lighter shadowed area corresponds to the [5 95] percentiles of the distribution of the analytical estimator and the Monte Carlo estimator defined in Eq:10, which showed the same variance. Right Panel: scatter plot of the split-half (green) and the Monte Carlo (blue) noise estimators against the value of the analytical noise ceiling estimator. The scatter plot comprises the noise values for all the levels of experimental noise used at the left panel.

### Influence of sample size and the interaction with the regularization parameter

The left panel of Fig 4 shows, on simulated data, the performance (measured as the correlation between predicted ***β***\* and 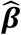 vectors) of the model which generated the data as a function of sample size. In addition, because when fitting linearized computational models to fMRI responses (see e.g [35]) the number of trials is in the order of the number of features (or an order of magnitude smaller) and the features show some degree of collinearity. Regularization (e.g. Ridge Regression) is typically used to alleviate these issues, and we consider here its influence on the performances with respect to the noise ceiling. The samples varied from the original *n* = 168 up to 10 times more the number of features *n* = 1680. The experimental noise was fixed to 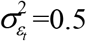. Using Ridge Regression predicted responses were obtained fitting the model used to generate the simulations and we report, for simplicity, the performances using two different regularization parameters λ = 10^0^ and λ = 10^3^ (green and red curves respectively in Fig 4).

**Fig 4.**
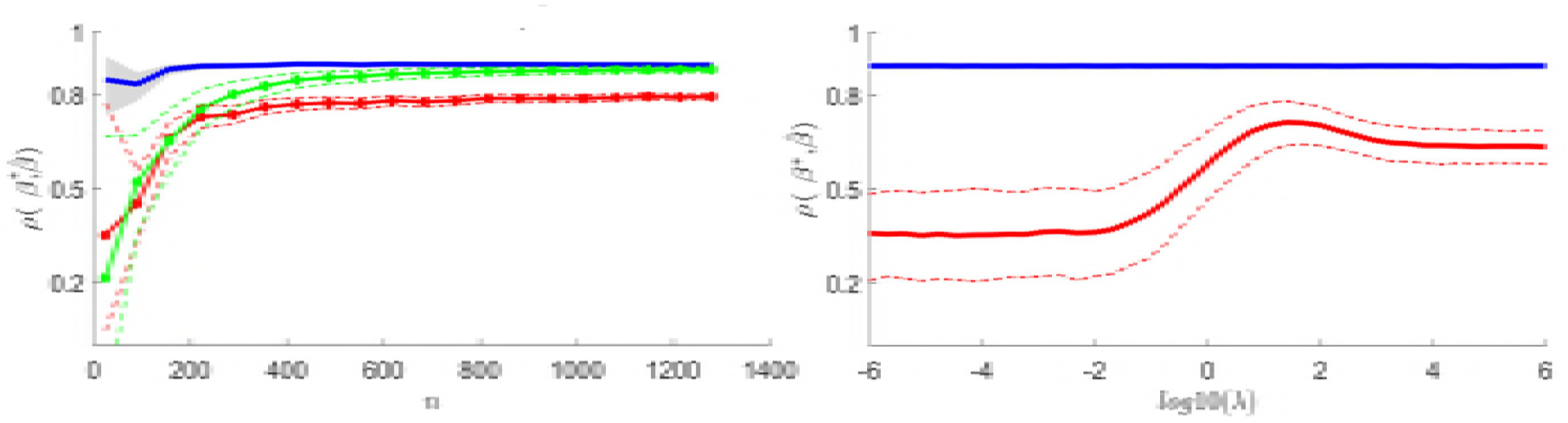
Left panel: Model performance as a function of the number of trials for a fixed level of experimental noise. The analytical noise ceiling estimator for the correlation coefficient in blue and its [5 95] percentiles denoted by the shadowed area is plotted together with the performance of two models that differ in the amount of regularization (green λ = 10^0^; red λ = 10). The corresponding [5 95] percentiles are presented in green and red dotted lines. Right panel: The effect of regularization on the performance. The correlation coefficient is used the metric for describing the performance. The blue line depicts the analytical noise ceiling estimator which is independent of *λ* The mean accuracy of the model performance and the [5 95] percentiles of the distribution across 1000 permutations are presented by the red dotted line.

As expected, prediction accuracy increased asymptotically with sample size. However, as a consequence of the bias-variance trade-off introduced by the regularization procedure, the performance did not reach the noise ceiling (despite of using the true generative model to predict the responses) even for a very large number of samples. High regularization implies less variability in the estimated linearized model weights (i.e. the pRF in fMRI encoding approaches) as depicted by the narrower [5 95] variability bands. Note that the reduced variability in data poor scenarios comes at the cost of an increased bias highlighted by the larger distance to the noise ceiling in data rich scenarios.

The right panel of Fig 4 depicts in more detail the influence of the regularization parameter. By keeping the experimental noise and sample size constant (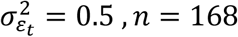 respectively) we evaluated the prediction accuracy (measured as correlation) of the model underlying the generation of the data and considered the effect of regularization on the difference between the actual model performance and the noise ceiling. The performance did not reach the noise ceiling even when the optimal regularization parameter is selected (the one that results in the highest performance).

### Testing the noise ceiling by introducing mismodelling at the level of the computational features

The observation that the use of regularization introduces a bias in the performance of the true generative model may not be unexpected but inspires some considerations. In particular, the scenario in which a model other than the one underlying the generation of the data is closer to the noise ceiling needs to be taken into account.

In the following we present the results obtained in different mismodelling scenarios, their effect on the performances of regularized linear models and their relation to the noise ceiling (here estimated using the proposed closed form estimator – Eq:13). In Fig 5 we report the results of an analysis evaluating the effect on performance of adding noise to the model underlying the generation of the data (without changing its complexity – i.e the number of columns of the computational model). We fitted the models using Ridge Regression and we report, for simplicity, the performances using λ = 10. We present variable levels of modelling error 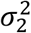, ranging from zero (no mismodelling green curve in Fig 5) to having the same variability that the deterministic part of the model: 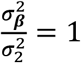 (gray region in Fig 5). As expected, when noise is added to the generative model the performance decreased below the performance of the model that generated the data (green curve). In line with the previous results, the use of regularization made all models fall short of the estimated noise ceiling.

**Fig 5.**
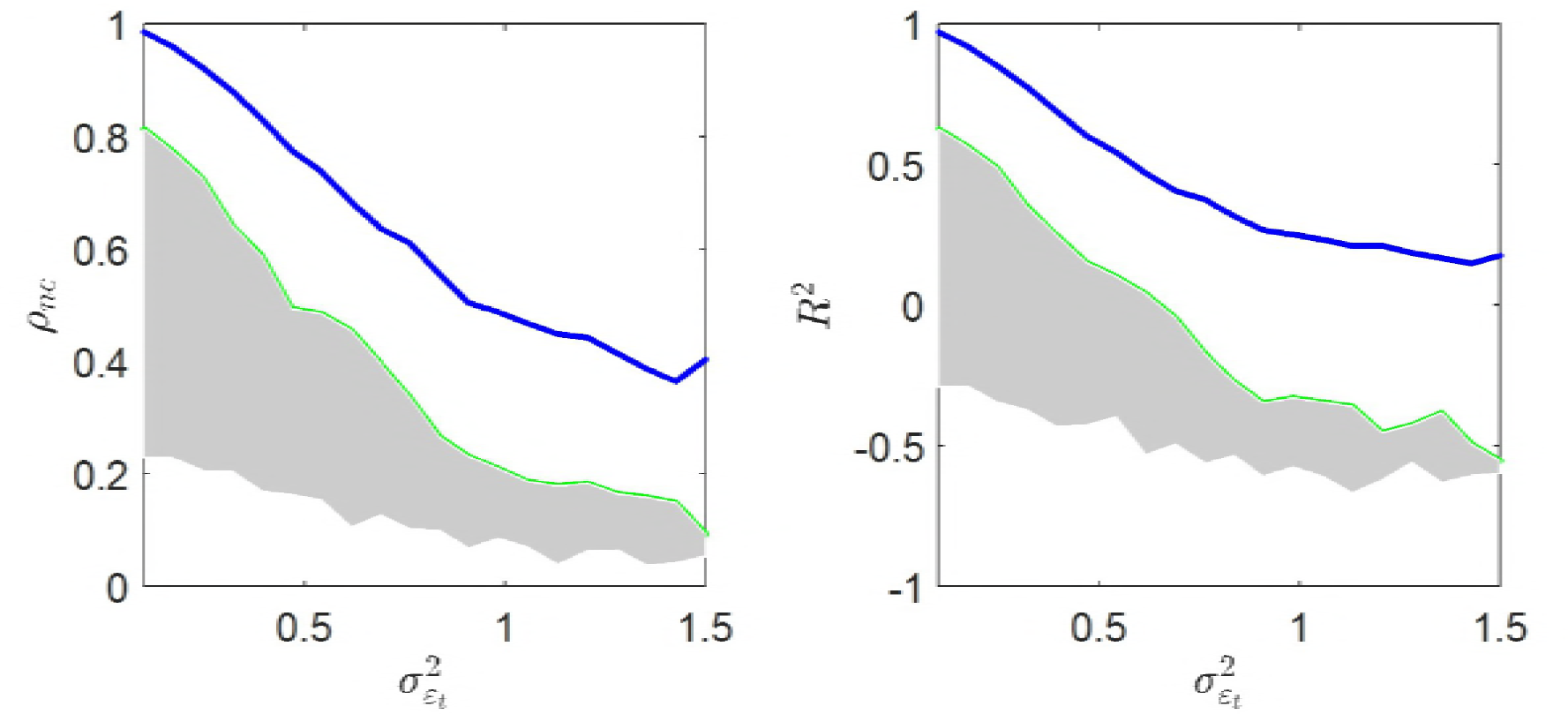
Influence of missmodelling effects on the performance of pRF models. The analytical noise ceiling estimator is denoted by the blue line as a function of the experimental noise. The shadowed area denotes the variation in model performance as a function of the variance of the *ε*_2_. The results are presented for the correlation coefficient (left panel) and explained variance (right panel).

### Influence of model complexity on the model performance

While adding noise to the model deteriorates its performance, we also wanted to consider the impact of model complexity. In simulations, we considered a realistic scenario (see e.g [7]) in which the model underlying the generation of the data is a model that represents sounds in the space of frequencies (120 logarithmically spaced values between 250 Hz and 5.2 KHz), temporal rates ([1, 3, 9, 27] Hz) and spectral scales ([0.5, 1, 2, 4] cyc/oct). Instead of adding noise to these 128 features (as in the results reported in Fig 5), we considered the performances of a “reduced” model in which the frequency information was resampled in 8 logarithmically spaced bins (the features corresponding to the temporal and spectral rates are identical between the reduced and the original model). This “reduced” model (16 features) explained 93% of the variance of the complete (128 dimensional) model. Fig 6 shows that the model underlying the generation of the data performed better than the reduced model when the experimental noise is low. When the experimental noise increases, small differences between computational models (7% of variance in this case is not shared between model features) were negligible.

**Fig 6.**
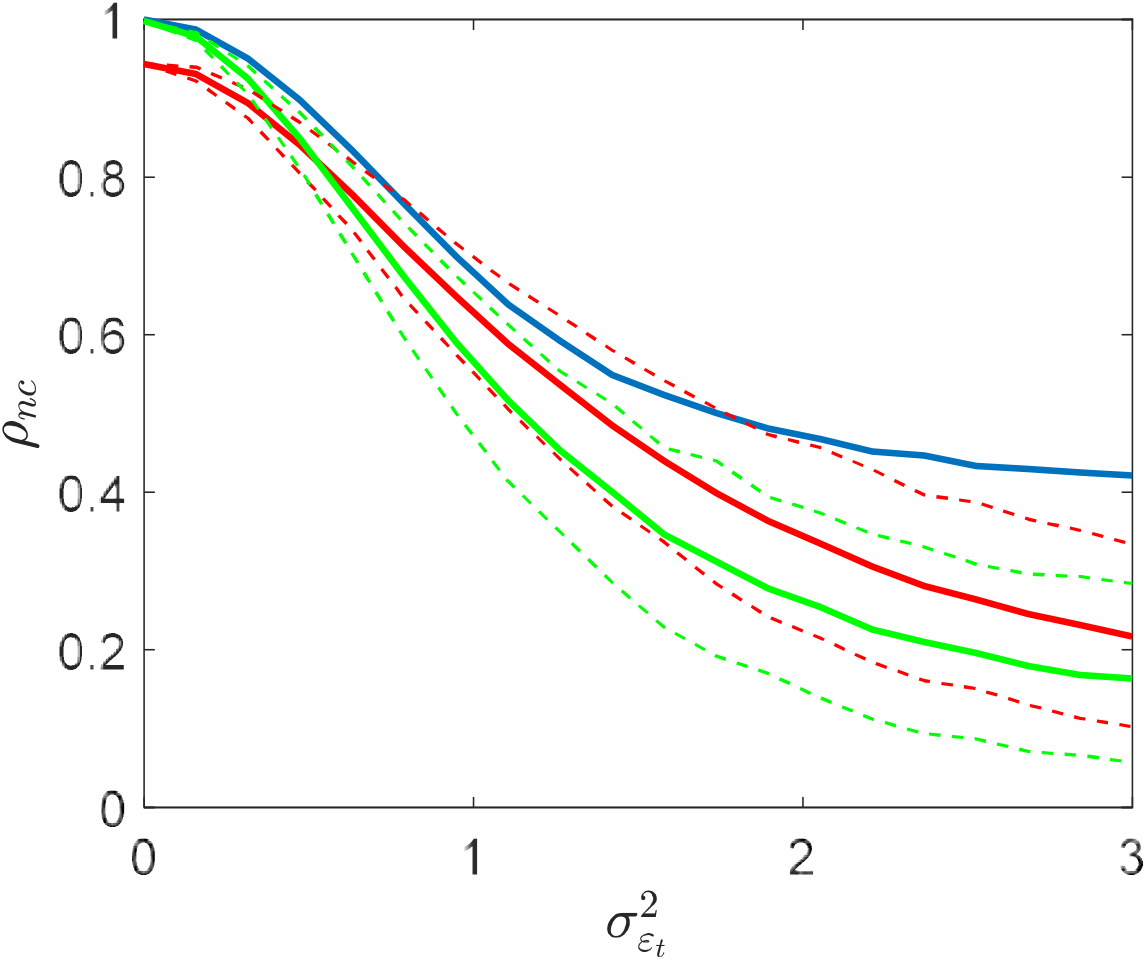
The figure shows how computational models of different complexities performed as a function of the experimental noise 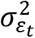. The data was generated with the f-128 model (green line) and evaluated used the f-128 model and the f-16 model (red line). The analytical noise ceiling estimator is depicted in blue.

In a second set of simulations we considered the effect of regularization by keeping constant the experimental noise. The regularization parameter was used as an indicator of model complexity. Fig 7 shows the performance of the model underlying the generation of the data (green curve) and reduced model (red curve) for different levels of regularization and in two different scenarios. The first (left panel in Fig 7) is the case in which the original (128) features have all the same importance for encoding the response, which implies that the scale of the coefficient of each feature of the model (i.e. the pRF) is the same. The results indicate that, in this scenario, at an optimal level of regularization the model underlying the generation of the data performed better than the reduced model. At the same time, the reduced model showed a lower dependence with respect to the regularization parameter. In the second set of simulations (right panel in Fig 7) we assumed frequencies and rate/scale to contribute equally to the responses by scaling the coefficients the features of the model in such a way that the coefficients for temporal rates and spectral scales have higher variance (120/8 times) compared to the coefficients for the frequencies. In this case and in the presence of experimental noise the reduced model outperformed the model underlying the generation of the data for all values of the regularization parameter.

**Fig 7.**
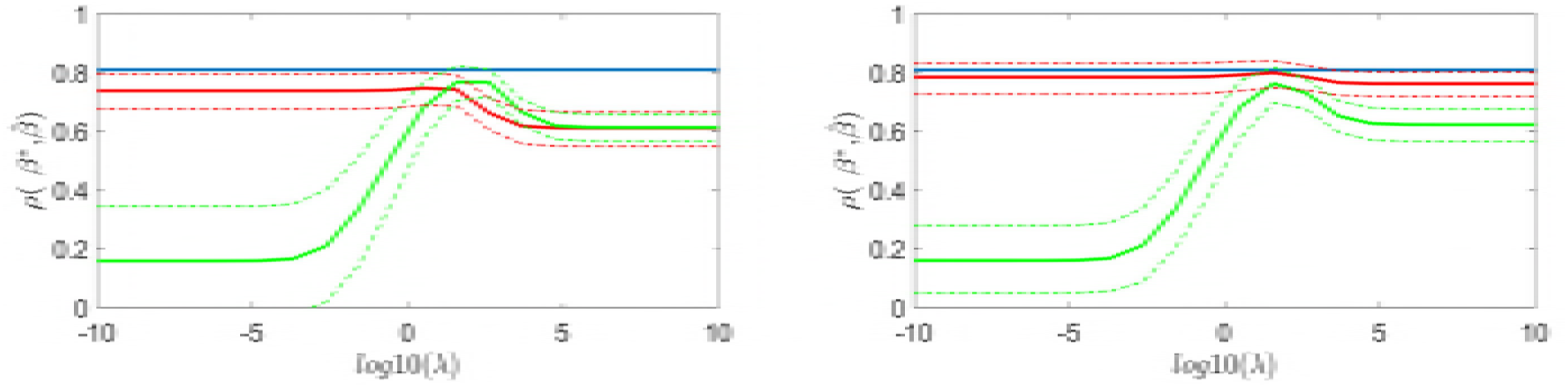
Influence of model complexity on the performance of pRF models. The left panel corresponds to the scenario were all features have the same importance. The right panel corresponds to the scenarios of two blocks of features of 8 and 120 with equal importance. The red lines shows the mean value and [5 95] percentiles of the performance of the f-8 model, while the green line shows the corresponding values for the f-128. The analytical noise ceiling estimator is depicted in blue.

### Noise Ceiling in a real data example

The analytical noise ceiling and the split-half noise ceiling were computed for cortical and subcortical voxels in three subjects. The MCnc estimator was not used in this analysis since it has the same expected value than the analytical noise ceiling and it requires larger computational burden. Initially the analytical noise ceiling was computed using the parametric estimate of the 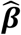 variances from OLS (see Eq. 6). Next, to evaluate the reliability of the OLS variances we also computed the noise ceiling using a non-parametric estimator of the 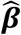 variances. Non-parametric estimates of the 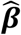 variance guarantee valid inferences even when the assumptions of the OLS/GLS model are violated [36]. In the case that OSL/GLS parametric assumptions are valid we should expect the parametric and non-parametric variances estimates to be similar and consequently the derived noise ceilings to be similar. The non-parametric estimate of the variance was computed with using block bootstrap randomization across the runs of the test set data [26]. The block bootstrap randomization preserves the original temporal dependencies of the fMRI noise within fMRI runs, assuming that the noise is independent between the fMRI runs. In practice the procedure works as follows: first the predicted values and the residuals of the fMRI time series are obtained using the OLS estimator. Next a new fMRI time series is constructed using the time series predicted with the OLS estimator with the addition of residuals that were resampled with replacement across runs. With the new fMRI time series the bootstrap based 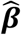 are obtained and their variance is computed across all the bootstrap samples. A total of 1000 bootstrap samples were used. The bootstrap based 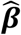 variances can be used as input to calculate the analytical noise ceiling in Eq. 13 (by replacing the variances estimated with OLS).

No significant differences were found between the cortical and subcortical voxels in the distribution of the noise ceiling values (Fig 8). In Table 1 we report the mean and variance for each noise ceiling estimator together with the correlation coefficients across voxels between them. The split-half estimator resulted in the lowest NC values in the three analysed subjects. The analytical noise ceiling based on the variances obtained with bootstrap resulted in estimates in between the SHnc and the analytical NC based on the variances obtained with OLS (parametric). Across all voxels the SHnc showed lower correlation with the analytical NC based on a parametric estimate of the variances than with the analytical NC based on bootstrapped variances.

**Fig 8.**
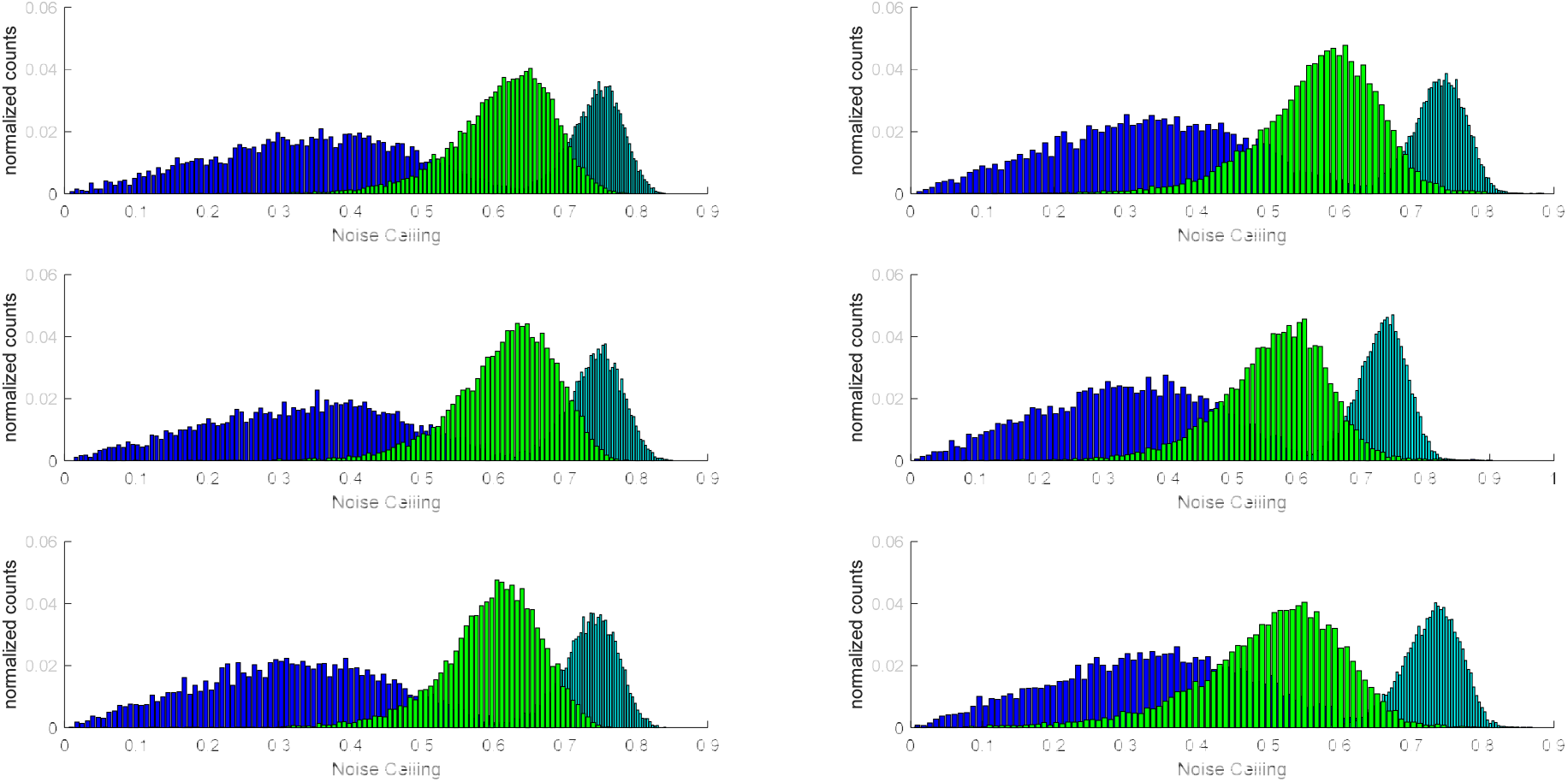
NC estimators for three subjects in the cortical (right column) and subcortical (left column) voxels. The normalized histograms of the NC across voxels is presented in blue for the split-half noise ceiling, green for bootstrap based analytical NC, and magenta for the parametric based analytical NC.

**Table 1.** The first three columns shows the mean and variance for the split-half noise ceiling, the bootstrap noise ceiling, and the noise ceiling based on the parametric variances for the voxels in the auditory cortex. The last three columns show the correlation between the noise ceiling estimators across voxels.

|  | Mean (variance) |  |  | Correlation between NC methods |  |  |
| --- | --- | --- | --- | --- | --- | --- |
|  | SHnc | Boot-nc | Param-nc | (SHnc,boot) | (SHnc,param) | (SHnc,boot) |
| S01 | 0.33<br>(0.02) | 0.57<br>(0.02) | 0.87<br>(0.0005) | 0.53 | 0.33 | 0.57 |
| S02 | 0.34<br>(0.02) | 0.56<br>(0.02) | 0.86<br>(0.0003) | 0.54 | 0.34 | 0.52 |
| S05 | 0.34<br>(0.02) | 0.51<br>(0.01) | 0.87<br>(0.0004) | 0.53 | 0.30 | 0.50 |

## Discussion

In neuroscience applications, computational models are a formal expression of the algorithms that underlie cognitive and sensory processes. Linking a model with non-invasive measures of brain activity (such as fMRI) [37, 38] allows testing the accuracy of the algorithm and eventually refine it to produce brain inspired models. Ultimately this approach can produce an understanding of brain function at the level of fundamental processing units. In this sense computational neuroimaging approaches represent a departure from classical subtraction based (hypothesis-testing) approaches (i.e. univariate GLM and multivariate decoding) [39] as they aim at understanding the response of a brain area to any given stimulus and do not focus (primarily) on the difference between (pre-defined) classes of stimuli. The accuracy of a model aimed at characterizing the function of a brain area is measured in terms of the ability to predict the responses (or representation) of these area to sensory (or cognitive) stimuli. Importantly, when reporting the results of an analysis based on computational neuroimaging techniques (e.g. linearized encoding, population receptive field modelling, representational similarity analysis), it is often recommended to report the performances (e.g. out of sample prediction in the case of fMRI encoding) with respect to the maximum performance that a “perfect” model would allow given the experimental variability in the data. In this sense, the intrinsic noise of the measurement procedure provides a bound to the performance of any model in predicting responses and this bound has been referred to as the noise ceiling. In some cases this recommendations have led to the use of normalized accuracy scores (e.g. dividing the accuracy by the noise ceiling, [11–15].

In this article, we evaluated two existing approaches for the calculation of the noise ceiling using simulations and real data. Additionally, assuming that modelling efforts are devoted to the prediction of the responses of single stimuli embedded in noisy (fMRI) time courses (e.g. distinguishing this approach from the otherwise used procedure of measuring the accuracy as the prediction of the whole times series), we proved that the noise ceiling can be computed based on the well-known property of variance partitioning of a two-level hierarchical model without the need of computationally demanding procedures. In simulations, we demonstrated that the previously introduced MCnc estimator [16] and the analytical estimator we derived here, have the same expected value and variance and as a consequence they have the same interpretation.

In real data, our analysis revealed a discrepancy between the SHnc and the analytical NC (or the MCnc) especially when the noise variance was estimated using a parametric model. The SHnc produced pessimistic values for the noise ceiling while the analytical NC with a parametric estimate of the variance of the noise produced apparently optimistic values for the NC. The SHnc is fully non parametric model that can account for violations of the OLS/GLS assumptions and random effects in the brain responses (e.g. across fMRI runs/sessions). Its calculation requires two independent estimates of the responses. In experiments that do not measure two independent (but identical in terms of presented stimuli) test sets, the SHnc can be calculated by splitting the available test data (i.e. by estimating the test set considering two independent partitions of the trials of each stimulus). A correction factor is applied to the SHnc to account for the reduced data. The analytical NC (and the MCnc) can produce optimistic values when the variance of the fMRI responses is underestimated as it is the case when the OLS (the most often used estimator for the responses in fMRI studies) assumptions are violated [40]. The results of the analysis of the real data show that the analytical NC (and as a consequence the MCnc) resulted in more conservative (and thus more accurate) estimates of the bound when the variance of the noise is calculated on the basis of a bootstrap procedure. This procedure allowed estimating a noise ceiling that is both robust to the violation of OLS assumptions (e.g. the presence of random effects) as well as not excessively conservative as in the case of the SHnc and represents, in our opinion, the appropriate method to use for the estimation of the noise ceiling in single fMRI voxels.

Computational models of sensory or cognitive process can consider a high number of parameters. In the case of linearized encoding models the stimuli representations in the parameters space is fitted to the voxels’ responses to a number of stimuli that, in some cases, can be one order of magnitude smaller. In these cases, regularization can be used to avoid overfitting, and the same procedure allows accounting for collinearity of the stimuli in the model parameter space. Importantly, it is a well-known property of regularization approaches the one to impose a bias-variance trade-off. In the context of comparing the performance of a computational model with the noise ceiling, the impact of the bias-variance trade off requires some consideration. Our simulations indicate that, when using regularization, the generative computational model (i.e. the model used to simulate the data and thus the best model to explain the data in a generative sense) cannot reach the noise ceiling. A relevant conclusion arises when analysing the dependence of the performance with model complexity. In noisy scenarios, a low dimensional model with a less precise representation of the stimuli (that captures only in part the variance of the generative representation) can outperform the generative model. This consideration can guide the selection of models to be fitted to the data and, as expected, draws attention to the dimensionality and the amount of shared variance of the models being compared in computational neuroimaging analyses.

It is important to stress that, the noise ceiling is itself a random variable with its corresponding variability. The expected value corresponds to the maximum performance having considered the noise in the data. However, for a particular (noisy) data sample (one test set) is possible that the performance of a model being tested is above the noise ceiling estimate on the same test set. Our results show that different estimators have different variability with the split half estimator exhibiting the larger variance, which is a second reason (together with the conservative nature of the SHnc estimator) for preferring an analytic (or Monte Carlo) estimate with bootstrapped variances. These considerations also grant a comment on the practice of reporting noise corrected performance values (see e.g. [41]. An evident advantage of this practice is that the noise corrected accuracies allow a direct comparison across experiments (or ROIs) with different levels of noise (e.g. acquired across imaging centers or changing MRI acquisition parameters). Nevertheless, considering that the noise ceiling itself is not an observable quantity that is estimated from the data with uncertainty (which is not independent of the mean), this practice is controversial. Different noise ceiling estimators have different variances and rely on different assumptions, and thus at the very least the exact procedure for the estimation of the noise ceiling should be specified. Reporting only noise ceiling corrected performances do not allow assessing (independently) the quality of the data and the observed effect size. Consider the following example: a voxel which reached an accuracy of 0.1 with an estimated NC of 0.2 would result in identical noise corrected accuracy 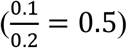 than a voxel which reached 0.4 with NC of 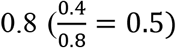. However the statistical significance and the effect size of an accuracy of 0.1 vs an accuracy of 0.4 differs in orders of magnitude and is computed on the uncorrected accuracies values since the null distribution of the noise normalized accuracies have not been determined yet. For these reasons, it seems more appropriate to report both the observed accuracies and the noise ceiling (see e.g. [10,42,43].

Here we focused on previous NC estimators that have been used for single fMRI voxels in single subject data. A related procedure, not included in this paper, is the noise ceiling developed for representational similarity analysis (RSA) [44] which is implemented in the RSA toolbox [45]. This approach estimates the noise ceiling for an RSA by considering the variability between subjects. The method consists in computing two bounds. The average correlation between each subject dissimilarity matrix and the mean dissimilarity matrix in the group provides the upper bound of the NC. The lower bound is computed using a leave one subject out procedure computing the correlation between the dissimilarity matrix of the left out subject and the mean dissimilarity matrix of the rest of the subjects. This procedure can be adapted to within subjects NC estimation and to single voxels instead of dissimilarity matrices (that are computed considering group of voxels). Depending on the experimental paradigm, brain responses can be estimated for every single run (if all stimuli are presented in all runs) or for every single trial. The two bounds can be computed (for each single voxel) considering: 1) the correlation between the response vector of a single run/trial with the average response vector across all trials/runs; 2) the correlation of the response vector a left out trial/run with the average response vector across the remaining trials/runs. We report the results of this procedure on our simulated data and in relation with the other NC estimators in the Appendix III (Fig 1). The results show that this approach produces lower NC values due to the higher variance of the single run/trial brain responses, note that when considering half of the runs/trials to estimate one response vector this procedure approaches the SHnc.

The framework proposed in this article is based in a two-level procedure for the estimation of the weights of the encoding models. One conceptual advantage of this two-level approach is that the residual variance can be partitioned in the variance of the experimental noise (level I) and the residual variance after fitting the model (level II). However, other approaches directly fit the encoding model convolved with the haemodynamic response to the fMRI time series [3]. In these approaches, the unexplained variance could be due to missmodelling effect (i.e. incorrect selection of model features) or also due to problems in the estimation of the haemodynamic response (i.e. incorrect selection of the basis functions) [46], or an interaction of both factors. This interplay between the haemodynamic response and the model makes it difficult to assess the experimental variability of the fMRI time series and as consequence the noise ceiling. One solution would be to consider the residual variance after the removal of the hemodynamic effect (estimated together with the pRF model in this case) and compute the NC of the combined estimation using the analytical method Eq. 13. Alternatively, the SHnc can be readily applied to the calculation of the NC at the level of the whole fMRI time series, but as we have indicated above, it may provide a conservative estimate.

## Appendix I: Relationship between the correlation coefficient and R2

The explained variance, under the assumption that the observations were normalized to have zero mean 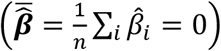 is:

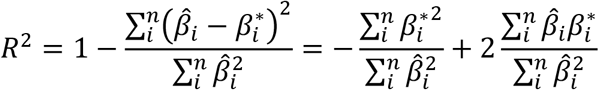

The correlation coefficient is under 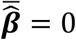 is:

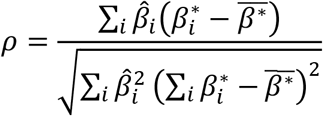

Writing *R*^2^ as a function of ρ produces

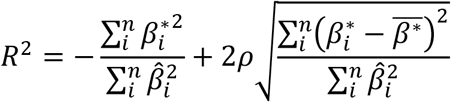

Based on the following the property of the unbiased estimator of the variance: 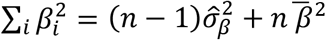 the relation between *R*^2^ and *ϕ* becomes:

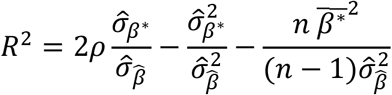

The assumption of using responses that are centered (normalized to have zero mean 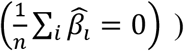 reflects the interest of encoding models in describing the variations in the response around their mean value as a function of the differences between the features of the presented stimulus. In the more general scenario where 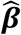 is non-centered the term 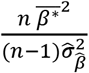 becomes into 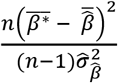 in the previous formula.

## Appendix II: Derivation of the analytical noise ceiling

The definition of noise ceiling for a vector of responses 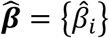 and vector of responses of the generative model ***β*** = {*β_i_*{ (not directly observed from the data) is (Eq. 11):

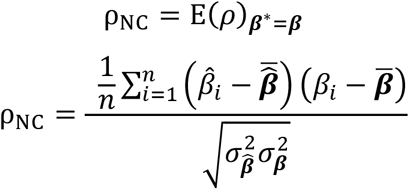

This definition refers to the covariation across the components *i* of the “noise free response” vector ***β*** and the estimated response 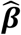 (noise contaminated). Note that under the generative framework, each component of the 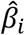 can be written as: 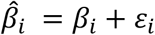, where *ε_i_* characterizes to the variability of each component of 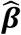 around the expected value due to experimental noise. The variance of *ε_i_* derived from the OLS is 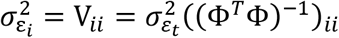, where the index *ii* refers to the diagonal entries of the corresponding matrix (see Eq:6). Substituting 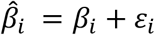 into the NC definition produces:

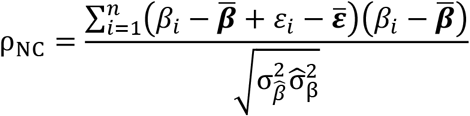

Noticing that the expected value of the estimation error each *ε_i_* is 0, and that *ε_i_* and *β_i_* are independent, the expected value of the noise ceiling becomes:

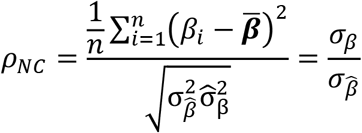

The estimator of *ρ_NC_* is obtained by substituting *σ_β_* and 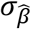 by its corresponding estimators.

The estimator of 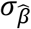 is 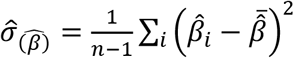. Consider that if 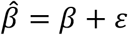 and *β* and *ε* are independent, then: 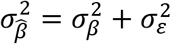. The estimator of 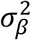 is:

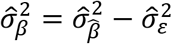

Where 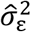 can be extracted from the OLS/GLS estimators of the variances of the 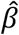 responses: 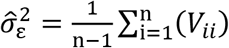. Substituting the estimators in the noise ceiling formula we obtain that the estimator of the noise ceiling is:

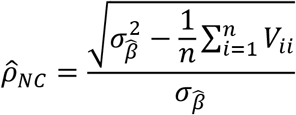

## Appendix III Adaptation of the RSA noise ceiling [45] for evaluating the performance of encoding models at individual subject level

The simulation consists in 24 identical fMRI runs of length 150 volumes each. At each fMRI run 20 stimuli were presented and each stimulus was repeated 3 times. The presentation onsets were linearly separated for across stimuli and then convolved with the canonical hemodynamic response. The brain responses used for generating the fMRI time series were sampled from the standard normal distribution and variable levels of experimental noise was added. The noise ceiling upper bound was obtained by averaging the correlations of the estimated responses at each fMRI run with the responses obtained with all runs together. The noise ceiling lower bound was obtained using a “leave run subject out” scheme. The brain responses for the run left out was correlated with the brain responses of the rest of the fMRI run combined. The correlations were averaged after transforming to the normal scale and then back projected to the correlation domain.

**Fig 1 (appendix III):**
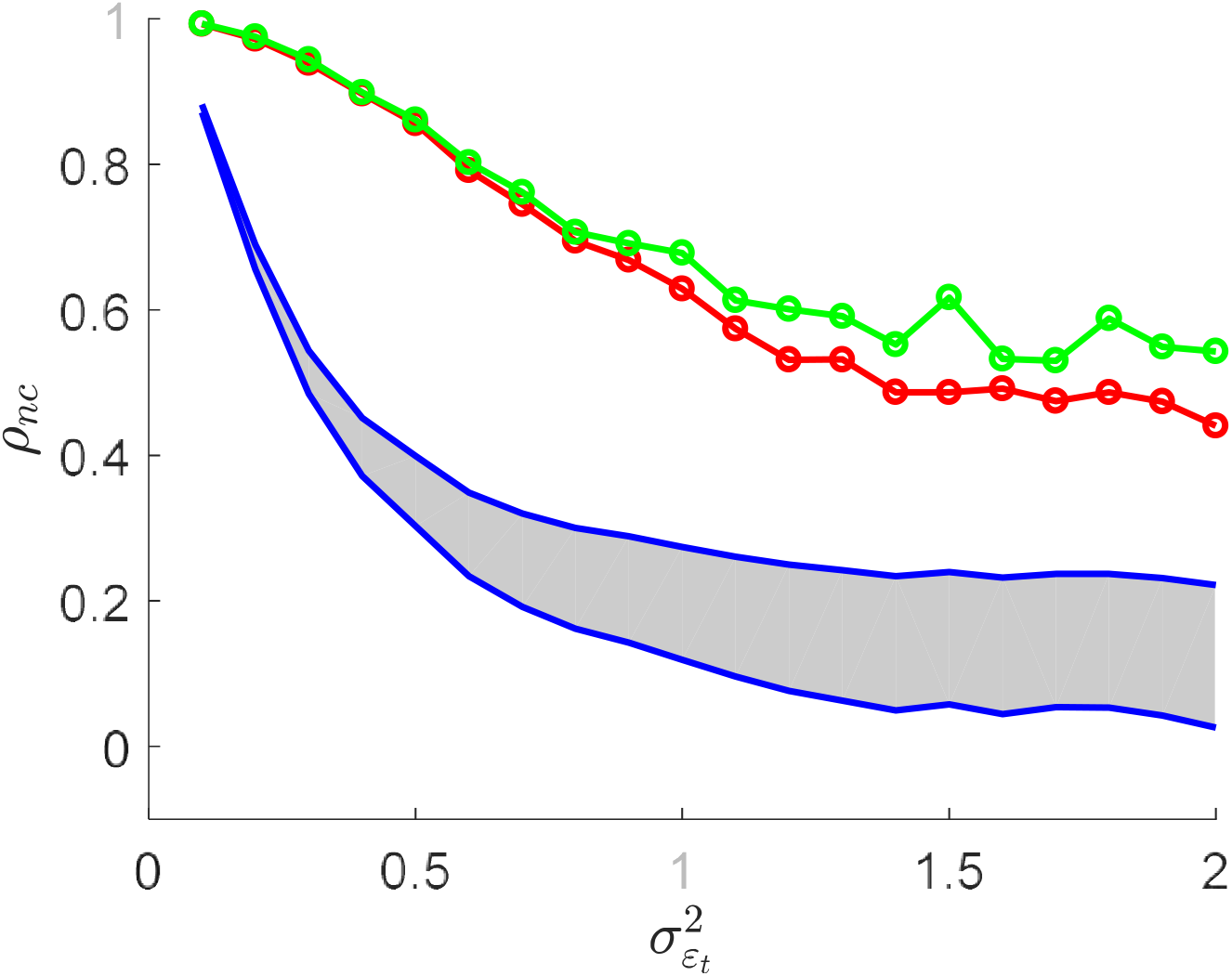
Comparison between the adapted RSAnc with the SHnc and the analytical noise ceiling for varying levels of experimental noise. The lower and upper bound of the RSAnc are denoted by the shaded region. The SHnc and the analytical NC are denoted by the green and blue curves respectively.

